# A Multiscale and Comparative Model for Receptor Binding of 2019 Novel Coronavirus and the Implication of its Life Cycle in Host Cells

**DOI:** 10.1101/2020.02.20.958272

**Authors:** Zhaoqian Su, Yinghao Wu

## Abstract

The respiratory syndrome caused by a new type of coronavirus has been emerging from China and caused more than one million death globally since December 2019. This new virus, called severe acute respiratory syndrome coronavirus 2 (SARS-CoV-2) uses the same receptor called Angiotensin-converting enzyme 2 (ACE2) to attack humans as the coronavirus that caused the severe acute respiratory syndrome (SARS) seventeen years ago. Both viruses recognize ACE2 through the spike proteins (S-protein) on their surfaces. It was found that the S-protein from the SARS coronavirus (SARS-CoV) bind stronger to ACE2 than SARS-CoV-2. However, function of a bio-system is often under kinetic, rather than thermodynamic, control. To address this issue, we constructed a structural model for complex formed between ACE2 and the S-protein from SARS-CoV-2, so that the rate of their association can be estimated and compared with the binding of S-protein from SARS-CoV by a multiscale simulation method. Our simulation results suggest that the association of new virus to the receptor is slower than SARS, which is consistent with the experimental data obtained very recently. We further integrated this difference of association rate between virus and receptor into a mathematical model which describes the life cycle of virus in host cells and its interplay with the innate immune system. Interestingly, we found that the slower association between virus and receptor can result in longer incubation period, while still maintaining a relatively higher level of viral concentration in human body. Our computational study therefore provides, from the molecular level, one possible explanation that this new pandemic by far spread much faster than SARS.

## Introduction

The coronavirus disease 2019 (COVID-19) has emerged at the end of year 2019 from Wuhan, a city in China, as a new pandemic [1, 2]. It has been found that the disease is caused by a new member of coronavirus family, SARS-CoV-2 [3, 4]. Like others in the same virus family, such as the coronavirus causing SARS [5, 6], SARS-CoV-2 is a single and positive-stranded RNA virus enveloped by lipid bilayer. The virus can capture and enter host cells in human through targeting specific receptors on their surface. Upon entry, the viral genome is released through membrane fusion. The released RNA genome of the virus is then replicated and translated into various types of viral proteins. The replicated RNA genome and synthesized viral proteins are finally assembled together into new viruses, before they escape and attack other cells [7, 8]. As a result, the infections of SARS-CoV-2 normally come with the similar symptoms as SARS, including fever, respiratory difficulty and pneumonia [9, 10]. Different from SARS, however, the new COVID-19 seems to have longer incubation period and thus is more contagious [11]. The disease has caused more than 43 million confirmed cases with at least 1 million death globally, according to the data from the World Health Organization (WHO) on October 27^th^ 2020. Therefore, the development of vaccine or therapeutic treatment for this ongoing public health crisis is highly demanding [12, 13].

Almost all the coronaviruses recognize their host cells through spike (S) proteins [14, 15]. S-protein is a glycoprotein expressed on the surface of viral envelop as homo-trimers [16]. Each S-protein further consists of two subunits. The S1 subunit includes a region called receptor-binding domain (RBD) which is used to target receptors in host cells, while the S2 subunit regulates the membrane fusion between virus and host cells [17]. These roles of S protein suggest that it could be a key target for vaccine and therapeutics developed to neutralize virus infection by blocking their invasion [18]. Moreover, it has been confirmed in a recent report that the new virus SARS-CoV-2 uses the same cell entry receptor ACE2 as SARS coronavirus [19]. The atomic structures of complex between human ACE2 and the RBD regions from S-protein of SARS-CoV have been obtained by x-ray crystallography [17]. It was also shown that the sequence of S-protein from SARS-CoV shares more than 70% identity with the S-protein from SARS-CoV-2 [6]. Therefore, it is reasonable to hypothesize that the new coronavirus uses the similar binding interface with ACE2 as SARS to enter host cells of human. The obvious follow-up questions are: whether the 30% variations in sequences between S-protein of SARS-CoV-2 and SARS can cause any difference in their binding to ACE2? Moreover, does this difference lead into any functional impacts on the life cycles of these viruses in host cells?

Comparing with the time-consuming and labor-expensive experimental studies, computational modeling serves as an ideal alternative approach to carry out fast tests on biological systems under the conditions that are currently inaccessible in the laboratory [20–26]. Therefore, here we developed a multiscale computational strategy to compare the process of recognition between the SARS-CoV and host cells with the interactions between the new coronavirus and host cells. A mesoscale model is used to simulate the process in which the coronaviruses are captured by ACE2 receptors on cell surface. We further constructed a structural model for complex formed between ACE2 and RBD of SARS-CoV-2 S-protein, so that the rate of their association can be estimated by a coarse-grained Monte-Carlo simulation and further compared with the binding of S-protein from SARS-CoV. Our simulation indicates that association of the new virus to the receptor is slower than SARS, which is consistent with the experimental data obtained very recently. We integrated this difference of association rate between virus and receptor into a simple mathematical model which describes the life cycle of virus in host cells and its interplay with the innate immune system. Interestingly, we found that the slower association between virus and receptor can result in longer incubation period, while still maintaining a relatively higher level of viral concentration in human body. Our computational study therefore explains, from the molecular level, why the new COVID-19 disease is by far more contagious than SARS. In summary, this multiscale model serves as a useful addition to current understanding for the spread of coronaviruses and related infectious agents.

## Results and Discussions

A rigid-body (RB) based model is first constructed to simulate the kinetic process about how viruses are captured by the cell surface receptors on plasma membrane. In brief, within a three-dimensional simulation box, the plasma membrane is represented by a flat surface below the extracellular region. The area of the square is 1 μm^2^, while the height of the simulation box is 500 nm. A number of ACE2 receptors (200) are initially placed on the membrane surface (pink in **Figure 1a**). They are represented by rigid bodies of cylinders and their binding sites are located on top of the cylinders (red dots in **Figure 1a**). The height of each cylinder is 10nm and its radius is 5nm. On the other hand, space above the plasma membrane represents the extracellular region. A number of coronaviruses are located in this area (golden in **Figure 1a**). Each virus is simplified as a spherical rigid body with a given radius (40nm). Trimeric S-proteins are uniformly distributed on the spherical surface of each virus (green dots in **Figure 1a**). Each S-protein can interact with an ACE2 receptor on plasma membrane. After any S-protein on one virus forms an encounter complex with a receptor, we assume that the host cell is captured by the virus. The dissociation between the virus and the receptor is not considered in the system, because we also assume that, after the association between S-protein and ACE2, the virus can enter the cell through membrane fusion. Following the initial random configuration, the diffusion of receptors and viruses, as well as their association, were simulated by a diffusion-reaction algorithm until the system reached equilibrium. The detail process of the simulation is specified in the **Methods**.

**Figure 1:**
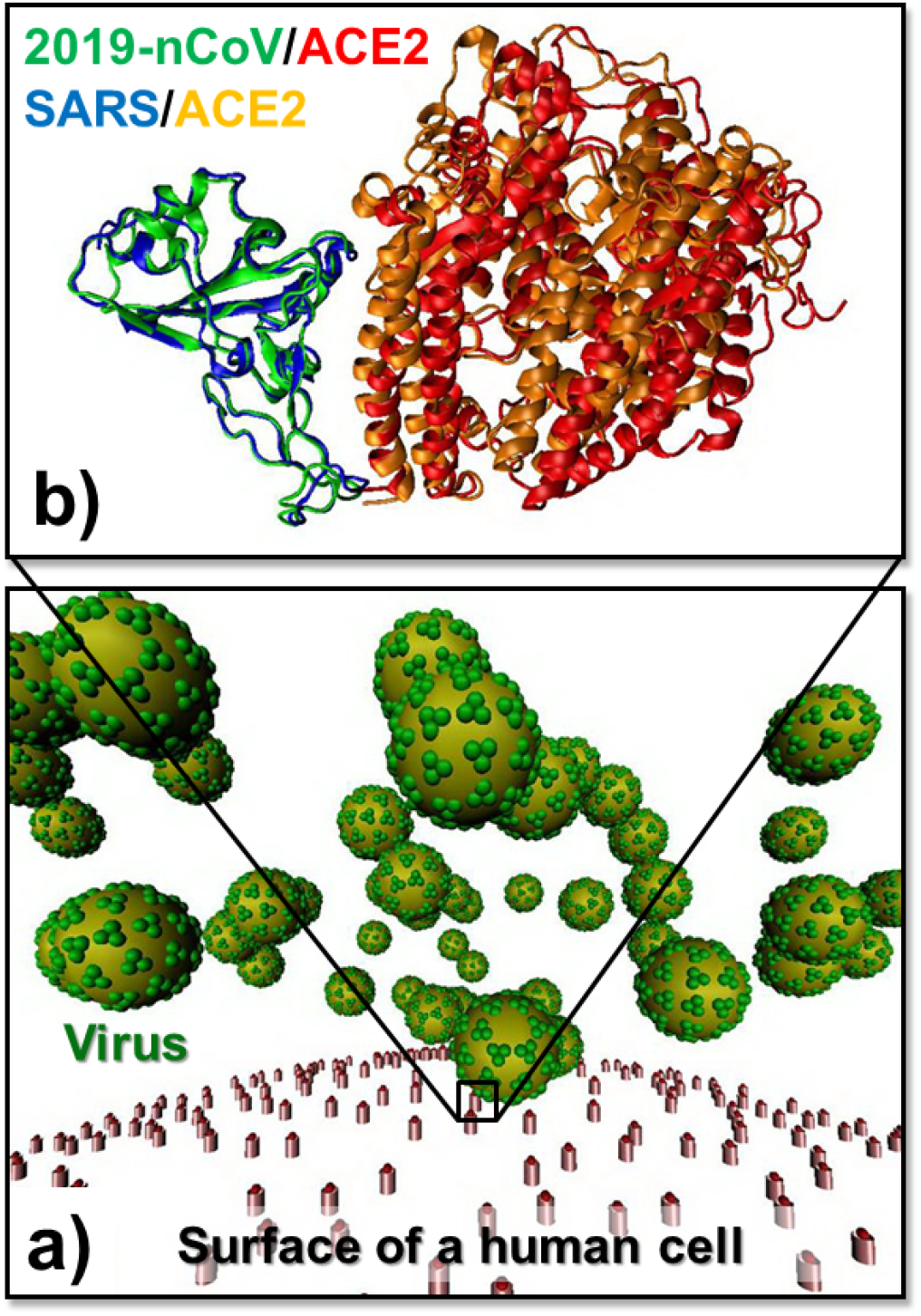
A mesoscopic model is constructed to simulate the kinetic process about how viruses are captured by the cell surface receptors on plasma membrane **(a)**. The plasma membrane is represented by a flat surface below the extracellular region, which contains a number of ACE2 receptors (pink). The space above the plasma membrane represents the extracellular region which contains a number of coronaviruses (golden). Each virus is simplified as a spherical rigid body with trimeric S-proteins uniformly distributed on its surface (green dots). Each S-protein monomer can interact with an ACE2 receptor on the plasma membrane. The rate of their association was estimated by Monte-Carlo simulations based on the structure models of complexes between ACE2 and RBD domains from S-proteins of different coronavirus **(b)**. The proteins in the figure are coded by different color index.

Before the rigid-body simulation, in order to provide a more realistic estimation on the binding between ACE2 and different coronaviruses, we specifically compared the S-protein from SARS-CoV-2 with the S-protein from SARS. We applied our previously developed residue-based kinetic Monte-Carlo (KMC) method to simulate the associate processes of these two systems. In detail, the atomic coordinates of the complex between human ACE2 and the RBD domain from the S-protein of SARS are taken from the PDB id 2AJF [17]. In parallel, the structural model of the complex between human ACE2 and the RBD domain from the S-protein of SARS-CoV-2 was computationally constructed, following the procedure described in the **Methods**. The structural comparison of these two protein complexes is shown in **Figure 1b**. For both systems, 500 trajectories based on their complex structures were generated by the KMC simulation which algorithm is specified in the **Methods**. In the initial conformation of each trajectory, S-protein and receptor were separated and placed with a random position relative to each other in which the distance between their binding interfaces is fallen within a given cutoff value *d_c_* of 20 Å. At the end of each trajectory, receptor and viral protein either form an encounter complex through their binding interface observed in the complex structure, or diffuse away from each other. Based on the simulation results collected from all the 500 trajectories, we counted how many times an encounter complex can be formed by the end of the simulation time, which gives the probability of association. As a result, the comparison of calculated probabilities of association for both systems is plotted in **Figure 2a**.

**Figure 2:**
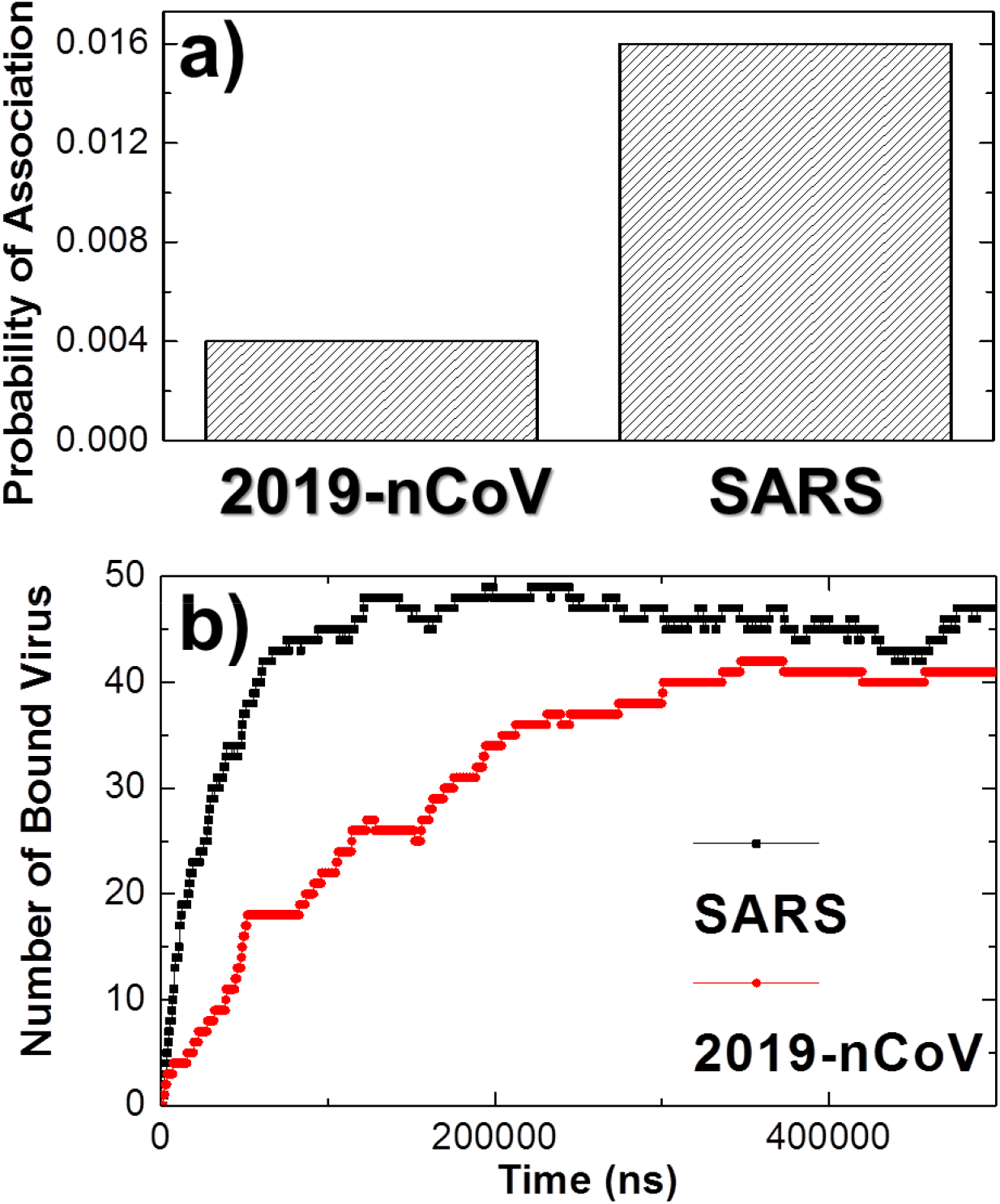
We used structure-based kinetic Monte-Carlo simulations to estimate the association between ACE2 and S-proteins from SARS-CoV and SARS-CoV-2. For each system, 200 simulation trajectories were generated. Based on these trajectories, the calculated probabilities of association are plotted in **(a)**. We found that the association between ACE2 and SARS-CoV S-proteins (right column) is faster than the association between ACE2 and SARS-CoV-2 S-protein (left column). We then fed the information into the rigid-body-based simulations. The simulations show the total numbers of viruses that were captured by host cells increased as a function of simulation time **(b)**. Moreover, we found that during the early stage of simulations, more SARS-CoV (black curve) attach to host cells than SARS-CoV-2 (red curve).

The figure shows that probability of association between ACE2 and the S-protein from SARS-CoV-2 is remarkably lower than the probability of association between ACE2 and the S-protein from SARS. Specifically, among the 500 simulation trajectories of SARS system, we found that 8 of them successfully formed encounter complexes, while among the 500 simulation trajectories of SARS-CoV-2 system, only 2 of them successfully formed encounter complexes. This result suggests that the association of the S-protein from SARS to the receptor is about four times faster than the association of the S-protein from SARS-CoV-2. The different of association rate from our simulation, interestingly, is confirmed by the experimental data that was measured very recently by Dr. McLellan’s lab [27]. Using surface plasma resonance (SPR), they showed that the association rate *k*_*a*_ of binding between SARS-CoV-2 RBD domain and ACE2 equals 1.36×10^5^M^−1^s^−1^, while the rate between SARS RBD domain and ACE2 equals 3.62×10^5^M^−1^s^−1^. Therefore, the experimental evidence indicated that the association of the S-protein from SARS to the receptor is about three times faster than the association of the S-protein from SARS-CoV-2, which is quantitatively consistent with our simulation results.

We then fed the information derived from the structure-based simulations into the rigid-body model. Two specific simulation systems were compared. A relatively fast rate of association between receptors and S-proteins on viral surfaces was adopted in the first system to represent the binding process of SARS, while a relatively slow rate of association between receptors and S-proteins on viral surfaces was adopted in the second system to represent the binding process of SARS-CoV-2. All the other parameters such as diffusion constants and concentrations in both systems remain the same. As a result, the total numbers of viruses that were captured by host cells are plotted in **Figure 2b** as a function of simulation time. Without surprise, the figure shows that although almost all viruses were attached to the cell surfaces by the end of both simulations, the kinetic process in the SARS system is much faster than the SARS-CoV-2 system, which is resulted from the difference in the association rate between receptors and their corresponding S-proteins. This leads into the fact that during the early stage of simulations, more SARS viruses attach to host cells than SARS-CoV-2. For instance, when the simulations in both systems reached the first 10^5^ nanoseconds, there have already been more than 40 SARS viruses attached to the cell surfaces. In contrast, there were less than 20 viruses attached to the cell surfaces within the same amount of time in the SARS-CoV-2 system. Considering that the function of a bio-system is often under kinetic, rather than thermodynamic, control [28, 29], we suggest that this time-dependent behavior is biologically more relevant. In reality, not all the viruses have the opportunity to find their target receptors on host cells. Many of them will be recognized and removed by our innate immune system. Therefore, the capability of how fast a specific type of coronavirus can target its receptors is especially critical to the process of its invasion, as well as the follow-up stages in its life cycle.

In order to further explore the impacts of our rigid-body simulation results on the rest process of virus infection after they invade into host cells, we proposed a mathematical model to delineate a simplified version of viral life cycle including its replication and packaging in host cells, the inflammatory signaling pathways due to the detection of foreign pathogens, and the follow-up inflammatory responses which lead to the apoptosis of infected cells and the removal of viruses by the recruitment of immune cells. The model can be summarized by the diagram shown in **Figure 3**. Specifically, coronavirus [V] can turn healthy cells [H] into infected cell [In] by binding to the receptors ACE2 on their surfaces. The RNA genome [m] is then released from bound virus [Vb], and then translated into different viral proteins. These proteins and replicated RNA are further packaged together as [P] in cytoplasm and finally escape from infected cells. On the other side, in order to avoid viral spread, our innate immune system triggers inflammatory signaling pathways in the infected cells [30]. For instance, the viral RNAs can be detected by RIG-I-like receptors (RLRs) [31]. The RNA binding of RLRs receptors initiates the signaling cascade by interacting with the mitochondrial antiviral-signaling (MAVS) protein [32]. The aggregation of MAVS on the surface of mitochondria will trigger the NF-κB signaling pathway that turns on gene expression of specific cytokines [S] to stimulate the inflammatory responses [33, 34]. The inflammation of host organism leads to the apoptosis of infected cells and the removal of virus by recruited immune cells such as microphages. In summary, the change of concentration for each variable in above system can be described by following set of ordinary differential equations (ODE).

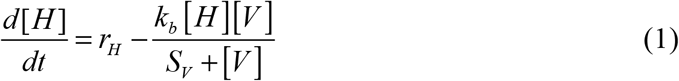

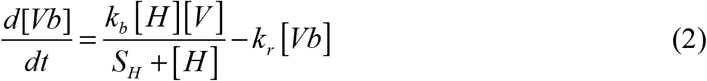

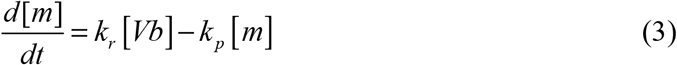

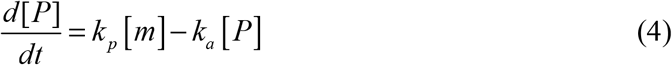

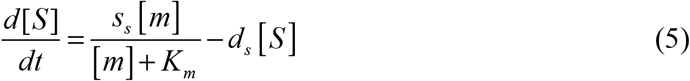

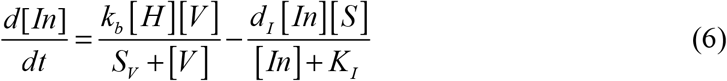

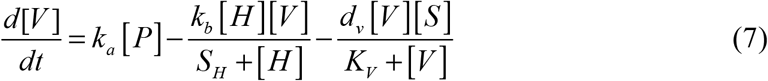

**Figure 3:**
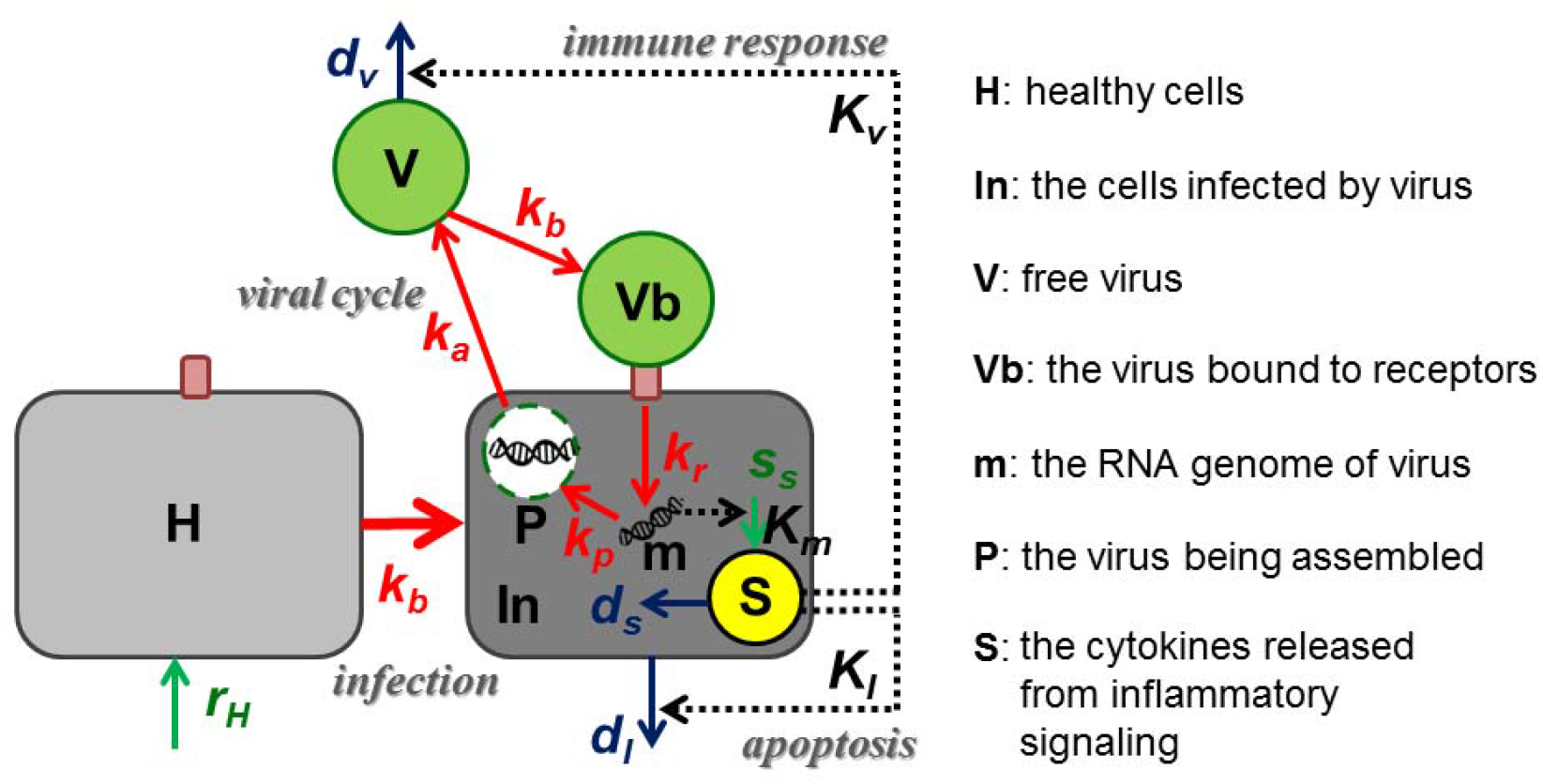
A mathematical model was proposed to delineate the simplified process of viral life cycle, including its replication and packaging in host cells, the inflammatory signaling pathways due to the detection of foreign pathogens, and the follow-up inflammatory responses which lead to the apoptosis of infected cells and the removal of viruses by the recruitment of immune cells. The meaning of each variable in the diagram is specified on the right. The system can further be described by a set of ordinary differential equations, as written from Equation (1) to Equation (7) in the main text.

Equation (1) describes the change of the healthy cells as a function of time in the system. The parameters *r*_*H*_, *k*_*b*_, and *S*_*V*_ in the equation represent cell generating rate, the rate and saturation coefficient of virus binding, respectively. Equations (2) to (4) indicate the entry, replication and packaging of virus in host cells, in which the parameters *k*_*r*_, *k*_*p*_, and *k*_*a*_ represent rates of viral protein translation, assembly and releasing. Equation (5) describes the stimulation of inflammatory signaling by viral RNA. The parameters *s*_*S*_, *K*_*m*_, and *d*_*S*_ in the equation represent the maximal activation rate, the saturation coefficient and the rate of degradation of inflammatory signals. Equation (6) gives how infected cells change in the system, while the parameters in the equation *d*_*I*_, and *K*_*I*_ indicate the rate and saturation coefficient of cell apoptosis that is stimulated by inflammatory signals. Finally, equation (7) suggests that the variation of total virus in the system depends on the release of newly packaged virus from infected cells, the binding of free virus to the healthy cells, and the immune response triggered by inflammatory signals. The parameters *d*_*V*_ and *K*_*V*_ in the equation give the maximal rate and saturation coefficient immune cells used to kill virus. Altogether, we solved above ODEs numerically by a stochastic simulation algorithm. The brief introduction of the simulation algorithm will be specified in the **Methods**.

Given predefined weights for all the simulation parameters and the initial values for each element, the dynamics of the system is evolved as a function of time. The simulation results of the mathematical model are summarized in **Figure 4**. As shown by the red curve in **Figure 4a**, the change of free virus in the system can be divided into three stages. It first decreases from its initial value. After reaching the minimal level, the number of virus then bounces back in the second phase until it drops again and finally vanishes at the end of the third phase. Relative to the free virus, the number of virus captured by host cells equals 0 at the beginning of the simulation. It increases in the first stage and diminishes in the second, as shown by the blue curve in the figure. In the third phase, very few viruses bound to host cells are detected in the system. Corresponding to the change of virus, the number of healthy cells is plotted as the black curve. The curve shows that the level of healthy cells reduces from the beginning and rises only after all free viruses are removed from the systems. Based on these kinetic profiles, the dynamics of the system can thus be interpreted as follows. In the first stage of simulation, free viruses invade into the healthy cells by binding to the receptors on their surface. We suggest that this stage corresponds to the incubation period. In the second stage, new viruses are assembled in and released from the infected cells. At the same moment, the viral genomes left in the infected cells stimulate the immune response. As a result, the total amount of virus is gradually lowered in the third phase, indicating that these viruses are cleared up by the innate immune system. After the removal of all viruses, the healthy cells in the system grow again, representing the recovery of the patient.

**Figure 4:**
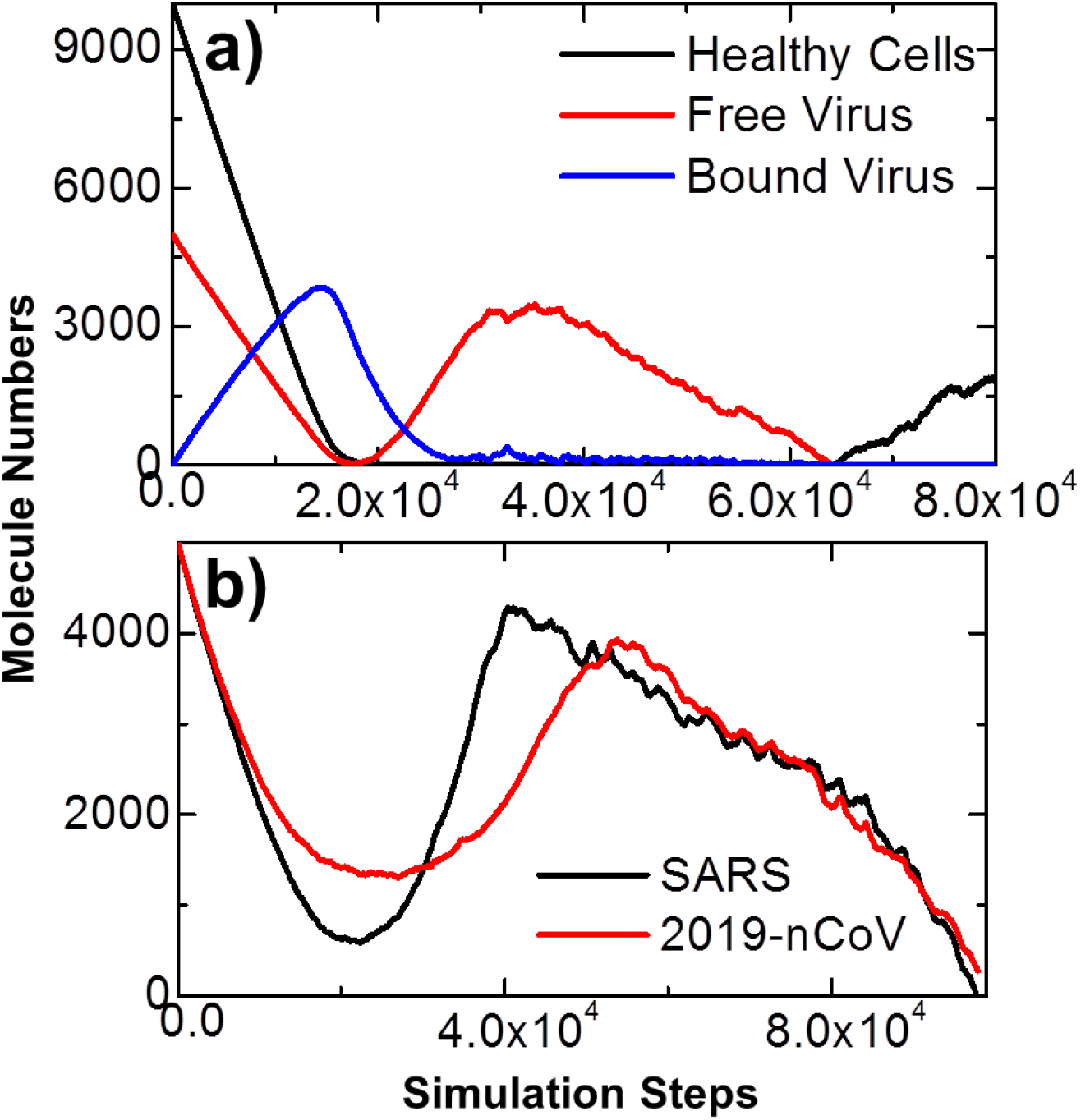
Given predefined weights for all the rate parameters and the initial values for each variable in the model, the dynamics of the system is evolved as a function of time by solving the mathematical model numerically with a stochastic simulation algorithm. The figure suggests that the dynamics of the system can be divided into three stages **(a)**. We further applied the model to two comparative systems. A relatively fast viral binding rate *k*_*b*_ was adopted in the first system to represent the binding process of SARS, while a relatively slow rate was adopted in the second to represent the binding process of SARS-CoV-2. As shown in **(b)**, we found that the first stage in the simulation of SARS-CoV-2 (red curve) is longer than the simulation of SARS (black curve), while the level of free virus at the end of the first period in SARS-CoV-2 is relatively higher than the corresponding level of free virus in SARS.

We further incorporated the results derived from the rigid-body simulations into the mathematical model to compare the viral life cycle in SARS and COVID-19. The rigid-body simulation suggests that SARS-CoV binds to receptor ACE2 faster than 19-nCoV, given the same amount of time. Therefore, we applied the mathematical model to two comparative systems. A relatively fast viral binding rate *k*_*b*_ was adopted in the first system to represent the binding process of SARS, while a relatively slow rate was adopted in the second to represent the binding process of SARS-CoV-2. All the other parameters such as diffusion constants and concentrations in both systems remain the same. As a result, the kinetic profiles indicating the changes of free virus level along with the simulation time in these two systems are shown in **Figure 4b**. The figure suggests that the first stage in the simulation of SARS-CoV-2 (red curve) is longer than the simulation of SARS (black curve). This is consistent with the clinical observation that the incubation period of COVID-19 could be as long as 14 days, while the incubation period is normally from 2 to 7 days. More interestingly, we found that the level of free virus at the end of the first period in the simulation of SARS-CoV-2 is relatively higher than the corresponding level of free virus in the simulation of SARS. Considering that a patient can contain a higher level of new coronavirus during his or her incubation period which is also longer than SARS, our simulation gives the insights about why COVID-19 is more contagious and spread faster than SARS [11]. Finally, when both simulations came to their third phases, we found that the total amount of virus of SARS is higher than SARS-CoV-2. This gives possible explanation about why the symptoms in most COVID-19 patients are relatively mild and not as severe as the symptoms in SARS patients [35].

In summary, it is important to point out that the life cycles of different coronaviruses can also be caused by many other factors such as the difference in the assembling pathways of these viruses. Additionally, the infectious response of each individual is also a case-dependent issue, relying on the expression levels of ACE2 receptors and the heterogeneity of immune response in different patients. Nevertheless, our multiscale computational model highlights the potential role of binding kinetics between human receptor and S-proteins from different coronavirus in, and provided the possible mechanism from the molecular level to their impacts on, regulating the dynamics of the entire viral life cycle.

## Conclusions

The recent outbreak of COVID-19 has drawn substantial attention especially after it spread to more than 200 countries and became a Public Health Emergency of International Concern (PHEIC) [2, 36, 37]. The pandemic is caused by a new type of positive-stranded RNA virus, known as SARS-CoV-2. Similar as the coronavirus that leads to SARS, it has been confirmed that the S-protein in SARS-CoV-2 also mediates its recognition with the human receptor ACE2. However, the differences of receptor binding in these two virus systems and their underlying implications are not well understood. It has been found that the kinetic aspect of binding between biomolecules in many biological systems is usually more important to their functions. Here, using computational structural prediction and coarse-grained simulations, we first have compared the association rate of binding between ACE2 and the S-protein from SARS-CoV with binding between ACE2 and the S-protein from SARS-CoV-2. Consistent with the experimental data obtained very recently, we found association of the new virus to the receptor is slower than SARS. We further interrogate the impact of this result on the difference in viral life cycle between SARS and COVID-19. By incorporating the information derived from coarse-grained simulations into a mathematical model, we found that the slower association between SARS-CoV-2 and ACE2 can result in longer incubation period, while still maintaining a relatively higher level of viral concentration in human body. This multiscale modeling framework, therefore, can offer one possible molecular mechanism to explain why this new infectious disease spreads much faster than SARS.

## Methods

### Construct the structural models for the complexes between receptors and viral S-proteins

The atomic structures of complex between ACE receptor and different viral S-proteins are needed for the simulations of their association. The structural models of these protein complexes were prepared as follows. The complex structure between human ACE2 and the RBD domain from the S-protein of SARS was determined by the x-ray crystallography experiment (PDB id 2AJF) [17]. On the other hand, the structural model of the complex between human ACE2 and the RBD domain from the S-protein of SARS-CoV-2 was constructed by computational modeling. In detail, the atomic structure of SARS-CoV-2 S-protein RBD domain was first predicted by I-TASSER [38] based on the newly released sequence of SARS-CoV-2 [39]. In parallel, the backbone model of the complex between human ACE2 and the RBD domain from SARS-CoV-2 S-protein was generated using the template-based structure prediction tool COTH [40, 41]. We then superimposed the predicted atomic coordinates of SARS-CoV-2 S-protein RBD domain onto its relative backbone position in the complex model, and also aligned the atomic coordinates of human ACE2 from the crystal structure onto the relative backbone position in the complex model. As a result, the structural comparison of these two protein complexes is plotted in **Figure 1b**.

### Estimate the association between ACE2 and S-protein by kinetic Monte-Carlo simulation

A previously developed kinetic Monte-Carlo algorithm [42] was used to simulate the association between S-protein and ACE2. A coarse-grained model of protein structures is used. Each residue in a protein is represented by its Cα atom and the representative center of its side-chain. The simulation starts from an initial conformation, in which two proteins in the complex are separated and placed randomly. Following the initial conformation, each protein diffuses randomly within one simulation step. A physics-based scoring function containing electrostatic interaction and hydrophobic effect is used to guide the diffusions of proteins during simulations. Based on the calculated energy, Metropolis criterion is applied to determine if the corresponding diffusional movements can be accepted or not. The simulation trajectory will be terminated if an encounter complex is successfully formed through the corresponding interface observed in the constructed model of native protein complexes. Otherwise, above simulation procedure will be repeated until it reached the maximal time duration. This method was applied to study the association between ACE2 and both S-proteins from the two virus systems. For each system, 500 trajectories are carried out. Each trajectory starts from a relatively different initial conformation, but the initial distances between the binding interfaces of S-proteins and receptors in all trajectories are below 20Å. The probabilities of association were then derived and compared based on counting how many encounter complexes formed among these trajectories in the two systems.

### Model the cellular attachment of coronavirus by rigid-body diffusion-reaction algorithm

As described in the **Results**, a rigid-body (RB) based model is constructed to simulate the binding between coronaviruses and cell surface receptors ACE2 on plasma membrane. Given the model representation and a randomly-generated initial configuration (**Figure 2a**), the dynamics of the system is evolved by following a diffusion-reaction algorithm [43–45]. Viruses or receptors are selected by random order for stochastic diffusion within each simulation time step. A virus is free to diffusion throughout the simulation box, while diffusions of membrane-bound receptors are confined to the plasma membrane. Periodic boundary conditions along both x and y directions are applied. Moreover, viruses are not allowed to move below the plasma membrane. If any virus moves beyond the top of the simulation box, it will be bounced back. The amplitude and probability of translational and rotational movements for viruses and receptors are determined by its corresponding diffusion constant. Association between viruses and receptors are followed after diffusions. Association is triggered if the distance between any S-protein in a virus and the binding site of a receptor is below a predetermined cutoff value. The probability to trigger the association is determined by the association rate, which was estimated by the kinetic Monte-Carlo method described in the previous section. Assuming that viruses can enter the cell through membrane fusion after they associate with ACE2, the dissociation between the virus and the receptor is not considered in the system. Finally, as above diffusion-reaction process iterates in both Cartesian and compositional spaces, the system will finally reach equilibrium.

### Solve the ordinary differential equations of viral life cycle by stochastic simulations

We use the stochastic simulation algorithm (SSA) developed by Gillespie to model the processes of biochemical reactions from equation (1) to equation (7) [46]. The algorithm starts from the initiation of time and populations of each species in the simulation system. Within each simulation step, the rates for all reactions are re-estimated by the given parameters and updated population of corresponding species. One of these reactions is then randomly selected based on the calculation of their relative weights. Finally, populations for the corresponding species are updated. The simulation moves forward to the next step by adding the system time with τ, in which τ is an exponential random variable with the average value equals the reciprocal of the summation for all the reactions. Above process is iterated so that the populations of each species in the system evolve along the simulation time. The values of all parameters in the simulation were chosen to be within the biologically meaningful range. The choice of these parameters does not qualitatively affect the general dynamic patterns of the system.

## Acknowledgement

This work was supported by the National Institutes of Health under Grant Numbers R01GM120238 and R01GM122804. The work is also partially supported by a start-up grant from Albert Einstein College of Medicine. Computational support was provided by Albert Einstein College of Medicine High Performance Computing Center.

## Author Contributions

Y.W. designed research; Z.S. and Y.W. performed research; Z.S. and Y.W. analyzed data; Y.W. wrote the paper.

